# Carbon monoxide utilisation by *Thermanaeromonas* species and description of *Thermobium azorense* gen. nov., sp. nov.

**DOI:** 10.64898/2026.07.10.736077

**Authors:** Anastasia Galani, Melissa Antony Venancius, Ben Tumulero, Detmer Sipkema, Diana Z. Sousa

**Author notes:** Corresponding author: Diana Z. Sousa.

## Abstract

Syngas fermentation by carbon monoxide (CO)-utilising acetogens offers a sustainable route for converting gasified waste materials into value-added chemicals. In this study, we isolated a novel thermophilic CO-utilising bacterium, strain AZ2, from marine hydrothermal sediment collected on the island of São Miguel, Azores, Portugal. Strain AZ2 is an obligately anaerobic, spore-forming bacterium. Average nucleotide identity (ANI; 78.4-86.7%) and digital DNA-DNA hybridization (dDDH; 23.4-32.5 %) analyses indicate that strain AZ2 represents a novel species within a previously uncharacterised lineage represented by the GTDB placeholder genus UBA2545 in the Neomoorellaceae family. Strain AZ2 was able to grow fermentatively on CO, producing acetate. We further demonstrated that its closest isolated relatives – *Thermanaeromonas toyohensis*, *T. burensis*, and *Thermanaeromonas* sp. strain 9S – are capable of growing on CO, producing either acetate or hydrogen gas (H_2_). Additionally, we unveiled the genomic potential for CO utilisation within other members of the GTDB placeholder class DSM-521 (previously Moorellia) to which our isolate belongs, expanding the list of possible thermophilic CO-utilising acetogens. We propose that strain AZ2^T^ represents the type strain of a novel genus and species, named *Thermobium azorense* gen. nov., sp. nov. (= DSM 121889^T^ = JCM 39698^T^).

## Introduction

Gasification of wastes, such as municipal or agricultural wastes, followed by microbial fermentation of the resulting syngas (CO, CO_2_, and H_2_), can address two environmental issues: it prevents environmental harm from waste and provides a sustainable production platform to meet commodity demand while reducing fossil fuel dependence (Liew *et al*. 2016b; Köpke and Simpson 2020; Fackler *et al*. 2021).

Syngas fermentation employs anaerobic microorganisms that utilise CO and/or CO_2_/H_2_ as carbon and energy sources. Carbon monoxide-utilising microorganisms, known as ‘carboxydotrophs’ (King and Weber 2007), have been isolated from different environments and belong to different taxonomic groups (Sokolova *et al*. 2005, 2007; Bae *et al*. 2006; Slepova *et al*. 2006; Slobodkina *et al*. 2022). They rely on nickel-iron-containing CO dehydrogenases (Ni-Fe CODHs), which can function either as monofunctional enzymes (CooS) or as bifunctional CODHs coupled with acetyl-CoA synthase (ACS). Monofunctional CODHs are primarily involved in CO oxidation. Depending on the microorganism, the final electron acceptor varies: protons in hydrogenogens (Sokolova *et al*. 2005; Yoneda *et al*. 2013; Fukuyama *et al*. 2019, 2020), CO_2_ in methanogens (Rother and Metcalf 2004; Diender *et al*. 2016) or acetogens (Drake and Daniel 2004; Bertsch and Müller 2015), and sulfate in sulfate-reducers (Parshina *et al*. 2005; Visser *et al*. 2014). The bifunctional CODH-ACS is a key enzyme in the Wood Ljungdahl pathway (WLP) used to fix two CO_2_ molecules into acetate (Svetlitchnyi *et al*. 2003; Ragsdale 2008; Schuchmann and Müller 2014). Within this complex, CODH catalyses the reduction of CO_2_ to CO, which is the actual precursor for the carboxyl group in acetyl-CoA. Notably, under in vitro conditions, CODH/ACS can also catalyse the oxidation of CO (Carlson, Papoutsakis and Kelly 2017).

Acetogens are promising biocatalysts for syngas fermentation, as they are able to use the WLP to convert CO or CO_2_ into acetyl-CoA, a precursor for various chemicals. So far, research has focused extensively on mesophilic carboxydotrophic acetogens, such as *Clostridium ljungdahlii, C. autoethanogenum, Eubacterium limosum,* or *Acetobacterium woodii* (Dykstra *et al*. 2022; Fernández-Blanco, Veiga and Kennes 2022; Rückel *et al*. 2022; Moon *et al*. 2024). Thermophilic acetogens remain underexplored, in part due to the limited availability of isolated strains. Acetogenesis from CO has been shown in thermophiles, such as *Neomoorella* species (Slobodkina *et al*. 2022; Harahap and Ahring 2023; Rosenbaum and Müller 2023; Böer *et al*. 2024; Galani *et al*. 2025), *Thermoanaerobacter kivui* (Weghoff and Müller 2016), *Aceticella thermautotrophica* (Frolov *et al*. 2023), and *Archaeoglobus fulgidus* (Henstra, Dijkema and Stams 2007). Thermophilic CO-utilising acetogens have some potential benefits for syngas fermentation, such as generally faster growth rates, reduced risk of microbial contamination and cost-effective product separation via bio-reactive distillation (Liew *et al*. 2016b; Kato *et al*. 2021). Therefore, efforts toward isolating and characterising more thermophilic CO-utilising microorganisms are essential for understanding and exploiting their biotechnological potential.

In this study, we report the isolation and characterisation of a novel thermophilic, CO-utilising bacterium, designated strain AZ2, from marine hydrothermal sediment collected on São Miguel, Azores, Portugal. Strain AZ2 represents the type species of a newly proposed genus, for which the name *Thermobium azorense* gen. nov., sp. nov. is proposed. Furthermore, we show that the closest isolated relatives of strain AZ2 – *Thermanaeromonas toyohensis*, *T*. *burensis* and *Thermanaeromonas* sp. Strain 9S are capable of CO utilisation, which had not been previously reported. Finally, we extended our analysis to other members of the GTDB placeholder DSM-521 class (formerly Moorellia), to which our novel strain belongs, and through comparative genomics, demonstrated that most members of this lineage possess the genomic potential for CO utilisation.

## Materials and methods

### Source of samples and microorganisms

A sediment sample was collected in September 2021 from Ponta de Ferraria on the island of São Miguel, Azores, Portugal (N 37°51.4984’, W 25°51.135’). At the time of collection, the sediment temperature was recorded at 60.2°C and the pH at 6.52. The sediment sample was collected in a 50 mL Falcon tube, which was filled to the top to minimise the presence of air. To ensure the preservation in anoxic conditions, the tube with the sample was stored in a vacuumed sealed bag along with Oxoid AnaeroGen^TM^ (ThermoFisher Scientific, Waltham, USA). The sample was kept at room temperature during transport to the lab and further processing.

Axenic cultures o*f Thermanaeromonas toyohensis* (DSM 14490^T^), *Thermanaeromonas burensis* (DSM 26576^T^) and *Thermanaeromonas* sp. strain 9S (DSM 22490) were purchased from the German Collection of Microorganisms and Cell Culture (DSMZ, Braunschweig, Germany) and were initially cultivated in accordance with the DSMZ instructions. Thereafter, the microorganisms were adapted to ‘AZ freshwater’ medium.

### Cultivation media

‘AZ marine’ medium contained (per L of medium): 0.14 g CaCl_2_*H_2_O, 0.33 g KCl, 0.14 g KH_2_PO_4_, 0.25 g NH_4_Cl, 0.5 mg NiCl_2_*6H_2_O, 0.5 mg Resazurin, 0.01 g (NH_4_)_2_Fe(SO_4_)_2_*6H_2_O, 6.5 g MgCl_2_*6H_2_O, 2.0 g NaHCO_3_, 19 g NaCl. For ‘AZ freshwater’ medium lower concentrations of NaCl (0.3 g L^-1^) and MgCl_2_*6H_2_O (0.1 g L^-1^) were used. Media were supplemented with 10 mL of trace element solution including (per L): 1.5 g N(CH_2_CO_2_H)_3_, 3 g MgSO_4_*7H_2_O, 0.5 g MnSO_4_*H_2_O, 0.1 g FeSO_4_*7H_2_O, 0.18 g CoSO_4_*7 H_2_O, 0.1 g CaCl_2_*2H_2_O, 0.18 g ZnSO_4_*7H_2_O, 0.01 g CuSO_4_* 5H_2_O, 0.02 g AlK(SO_4_)_2_*12H_2_O, 0.01 g H_3_BO_3_, 0.01 g Na_2_MoO_4_*2H_2_O, 0.03 g NiCl_2_*6H_2_O, 0.3 mg Na_2_SeO_3_*5H_2_O, 0.4 mg Na_2_WO_4_*2H_2_O (DSM medium 141). The medium was boiled and cooled on ice under continuous N_2_ flow. Unless stated otherwise, the pH was set to 7.0 using 1M NaOH. Afterwards, 47 mL of medium was dispensed into 117 mL bottles that were immediately capped with bromobutyl rubber stoppers (Rubber BV, Hilversum, the Netherlands) and aluminium caps. The headspace of the bottles was filled with the desired gas combination (e.g., H_2_/CO_2_/CO or N_2_/CO_2_/CO, to a final pressure ranging from 1.5 to 1.7 atm). Immediately after preparation, bottles were autoclaved at 121°C for 20 min.

Before inoculation, the medium was further supplemented with 1 mL L^-1^ of a filtered sterile vitamin stock solution containing (per L): 20 mg biotin, 200 mg nicotinamide, 100 mg para-aminobenzoic acid, 200 mg thiamine, 100 mg pantothenic acid, 500 mg pyridoxamine, 100 mg cyanocobalamin, 100 mg riboflavin (DSM medium 141). Finally, the medium was reduced with 1 g L^-1^ of L-Cysteine-HCl*H_2_O. When needed, substrates such as sugars, carboxylic acids and/or electron acceptors were added from sterile stock solutions to final desired concentrations.

### Enrichment culture and isolation of strain AZ2

To initiate the enrichment culture, 6% (w/v) of the sediment sample was inoculated into a 117 mL serum bottle containing 50 mL of marine AZ medium (initial pH 7.0) with a headspace of H_2_/CO_2_/CO (48:12:40 v/v) at 1.7 atm. Incubation was done at 60°C in the dark, without agitation. Upon growth, the culture was transferred (10% v/v, twice) in ‘AZ marine’ medium. At this stage, the culture was also transferred to ‘AZ freshwater’ medium, where faster growth was observed. Consequently, further enrichment was continued in the ‘AZ freshwater’ medium. After five subsequent 1% (v/v) transfers, a single morphotype was observed in the culture. The purity of the culture was determined by both phase contrast microscopy using a Leica DM2000 microscope (Leica, Microsystems, Weltzar, Germany) and direct Sanger sequencing of the full 16S rRNA gene (Macrogen, Amsterdam, the Netherlands). The Gram reaction of the culture was determined with a commercial Gram staining kit (Sigma-Aldrich, St. Luis, USA).

### Physiological characterisation of strain AZ2

Growth experiments were performed in ‘AZ freshwater’ medium with a headspace of N_2_/CO_2_/CO (48:12:40 v/v) at 1.7 atm. Cultures were incubated in the dark without agitation. To determine optimal growth conditions, strain AZ2 was cultivated at various temperatures (15-90°C), pH (5.0-9.5) and salinity (0-3%) values. The utilisation of different soluble (20 mM) and gaseous (1.7 atm) substrates was tested at pH 7.0 and 60°C. The substrates tested were: D-glucose, D-fructose, D-galactose, sucrose, xylose, maltose, pyruvate, butyrate, formate, propionate, succinate, lactate, malate, ethanol, methanol, H_2_/CO_2_ (80:20 v/v), N_2_/CO_2_/CO (48:12:40 v/v), CO (100% v/v). The utilisation of external electron acceptors was tested both with CO [1.7 atm N_2_/CO_2_/CO (48:12:40 v/v)] and with acetate (20 mM) as electron donors, at pH 7.0 and 60°C. The external electron acceptors tested were: nitrate (10 mM), sulfate (10 mM), thiosulfate (10 mM), fumarate (10 mM), perchlorate (10 mM), and Fe(III) oxide (20 mM). Soluble substrates and electron acceptors were added from sterile anoxic stock solutions before inoculation. Cultures were monitored by measuring the optical density of the cultures at 600 nm (OD600) and measuring substrate and product concentrations in the medium over time. When an external electron acceptor was added, both that compound and the corresponding reduced product were quantified. All tests assessing temperature, carbon sources, electron acceptors, pH, and NaCl tolerance were conducted in duplicate, whereas tests assessing growth on CO were performed in triplicate.

### Analytical techniques

Gaseous compounds (CO, H_2_ and CO_2_) were analysed with a gas chromatograph (Compact-GC 4.0; Global Analyser Solutions, Breda, The Netherlands) equipped with a thermal conductivity detector. A Carboxen 1010 column was used as a pre-column separating CO_2_ from other gases. A Molsieve 5A column, operating at 140°C, was used for measuring CO and H_2_, while an RT-Q-bond column, operated at 60°C, was used for measuring CO_2_. Argon was used as the carrier gas.

Organic acids, sugars and alcohols were determined using a high-pressure liquid chromatograph (LC-2030C, Shimadzu, Japan) equipped with a Shodex SH1821 column and UV and refractive index detectors. The system was operated at 40°C with a flow rate of 1 mL min^-1^. The eluent used was 0.01 N H_2_SO_4_. Sample preparation involved centrifugation of 1.0 mL samples for 15 min at 10,000 g at room temperature. The supernatant was then mixed with crotonic acid or arabinose (10 mM for both) as internal standards at a 2:3 (sample:internal standard, v/v) ratio.

Sulfate, thiosulfate, nitrate and perchlorate were measured with a ion chromatograph (ThermoICS2100, Dionex, Sunnyvale, CA, USA) equipped with a suppressed conductivity detector. For sulfate and thiosulfate, a Dionex AS17 column (250 mm x 2 mm) (Dionex, Sunnyvale, USA) set to a temperature of 30°C was used. For nitrate and perchlorate, a Dionex AS19 column (250 mm x 2 mm) (Dionex, Sunnyvale, USA) set to a temperature of 30°C was used. KOH was used as the eluent in a gradient, starting at 1 mM during the first four minutes and increasing to 40 mM by 20 minutes. The eluent flow rate was 0.3 mL/min for the Dionex AS17 column and 0.40 mL min^-1^ for the Dionex AS19 column. Samples (1 mL) were centrifuged for 15 min at 10,000 g at room temperature, and the supernatant was used for the analysis.

For peak integration and analysis, the Chromeleon™ data analysis software v7.2.9 (ThermoFisher Scientific, Waltham, MA), was used in all machines.

### 16S rRNA gene amplicon sequencing of the enrichment culture

DNA was extracted from the enrichment culture (after two subsequent 10% v/v dilutions) using the FastDNA™ Spin Kit for Soil (MP Biomedicals, Irvine, CA). 15 mL of culture was centrifuged in a 15 mL tube at 3,000 g for 15 min at room temperature. The supernatant was discarded, and 978 µL sodium phosphate buffer was added to the pellet. Homogenization of the mixture was done using a FastPrep™ 24 (MP Biomedicals, Irvine, USA) for 3 cycles of 60 s at a speed of 5.5 ms^-1^. For the DNA elution step, the catch tube was incubated at 55°C for 5 min before centrifuging the final eluate for increased elution efficiency.

PCR amplification was performed on the V4 region of the 16S ribosomal RNA (rRNA) gene with the 515F/806R primers (Parada, Needham and Fuhrman 2016). PCR was performed in triplicate as follows: 10 μL of 5 x green phusion buffer, 32.5 μL of nuclease-free water, 1 μL of dNTPs (500 μΜ), 0.5 μL of 515F primer (10 mΜ), 0.5 μL of 805R primer (10 mΜ), 0.5 μL of phusion hot start II DNA polymerase (2 U µL^-1^) and 5 μL of DNA template were mixed together for the PCR reaction. The PCR programme included 30 s of initial denaturation at 98°C and 30 amplification cycles, consisting of denaturation for 10 s at 98°C, annealing for 20 s at 47°C, and elongation for 60 s at 72°C. The final elongation was performed at 72°C for 3 min. The presence and length of PCR products were examined using electrophoresis in a 1% (w/v) agarose gel. Afterwards, the triplicate PCR products were pooled and purified with the HighPrep® PCR kit (MagBio Genomics, Alphen aan den Rijn, The Netherlands), according to the manufacturer’s instructions. The quality and quantity of the purified samples were examined with a NanoDrop (Thermofisher Scientific, Waltham, USA) and Qubit dsDNA BR assay (Thermofisher Scientific, Waltham, USA). 16S rRNA gene amplicon sequencing was performed with Illumina HiSeq, paired-end 150 bp (PE150) platform by Novogene (Beijing, China).

Processing of 16S rRNA gene amplicon data was done with NG-Tax v2.0 (Ramiro-Garcia *et al*. 2016; Poncheewin *et al*. 2020). The same tool was used for identifying amplicon sequence variants (ASVs) in the sample with Silva_138_SSU v138.1 as the reference database (Quast *et al*. 2013).

### Sequencing of the full 16S rRNA gene of strain AZ2

For sequencing the full 16S rRNA gene of strain AZ2, genomic DNA from a growing culture was extracted using the FastDNA™ Spin Kit for Soil, as described before. PCR amplification of the full 16S rRNA gene was performed using the 27F/1492R primer pair (Nübel *et al*. 1997). The PCR reaction mixture consisted of 10 μL of 5 x green phusion buffer, 36.5 μL of nuclease-free water, 1 μL of dNTPs (500 μΜ), 0.5 μL of 27F primer (10 mΜ), 0.5 μL of 1492R primer (10 mΜ), 0.5 μL of phusion hot start II DNA polymerase (2U/µL) and 1 μL of DNA template. The PCR program included 30 s of initial denaturation at 98°C and 25 amplification cycles, consisting of denaturation for 10 s at 98°C, annealing for 10 s at 50°C, and elongation for 10 s at 72°C. The final elongation was performed at 72°C for 7 min. The generated PCR product was examined with 1% (w/v) agarose gel electrophoresis and was purified with the DNA Clean & Concentrator™ – 5 kit (Zymo Research, Irvine, USA). Sanger sequencing with both the forward and the reverse primers was performed by Macrogen (Amsterdam, The Netherlands). The partial sequences were merged into a full-length sequence with Bioedit v7.2.5. (Hall 1990) and checked against the GenBank database (Sayers *et al*. 2020) using the NCBI BLASTn search tool (Altschul *et al*. 1990).

### Whole genome sequencing and analysis of strain AZ2, *Thermanaeromonas* sp. strain 9S (DSM 22490) and *Thermanaeromonas burensis* (DSM 26576^T^)

For whole genome sequencing, genomic DNA was extracted from a growing culture of strain AZ2 as described before. DNA from *Thermanaeomonas* sp. strain 9S (DSM 22490) and *T. burensis* (DSM 26576^T^) was obtained from DSMZ. Sequencing was performed with the Illumina NovaSeq 6000, paired-end 150 bp (PE150) platform by Novogene (Beijing, China). Initial inspection of the quality of raw reads was performed with FastQC v0.11.9 (Andrews 2010). The BBduk.sh script from BBtools v38.84 (Bushnell 2018) was employed for quality filtering and adapter removal (parameters: ktrim=r, k=23, mink=7, hdist=1, tpe, tbo, qtrim=rl, trimq=20, ftm=5, maq=20, minlen=50) as well as for the removal of sequencing artifacts and phiX contamination.

For the assembly of the quality-filtered reads SPAdes v3.15.2 (Prjibelski *et al*. 2020) was used, with the --isolate parameter enabled. Contigs shorter than 1000 bp were removed from the final assembly with the reformat.sh script from BBtools v38.84. The quality of the final assembly was determined with QUAST v5.0.2 (Gurevich et al., 2013). Mapping of the quality-filtered reads back to the contigs was performed with Bowtie2 v2.4.1 (Langmead and Salzberg 2012). The sequence alignment map (SAM) files, resulted from mapping, were converted into binary alignment map (BAM) files with SAMtools v1.18 (Danecek *et al*. 2021). The same tool was used for sorting and indexing the BAM files. Finally, the coverage of the genome was calculated with the ‘genomecov’ script from BedTools v2.29.1 (Quinlan and Hall 2010). Completeness and contamination values were calculated with CheckM v1.0.18 (Parks *et al*. 2015).

While analysing the genome of *T. burensis* (DSM 26576^T^) we detected contamination of the DNA. Therefore, contigs were binned with the binning module of metaWRAP v1.3.0 (Uritskiy, DiRuggiero and Taylor 2018) using three binning algorithms, as implemented in metaWRAP: metaBAT 2 (Kang *et al*. 2019), MaxBin 2.0 (Wu, Simmons and Singer 2016) and CONCOCT (Alneberg *et al*. 2014). Afterwards, the bin refinement module of metaWRAP was used to compare the bins and consolidate them into a final improved bin set. The metagenome assembled genome (MAG) representing *T. burensis* (DSM 26576^T^) was confirmed with taxonomic annotation using GTDB-Tk v2.4.0 (Chaumeil *et al*. 2020).

Initial annotation of the genome of strain AZ2 was performed using the Prokaryotic Genome Annotation Pipeline (PGAP v6.6) (Tatusova *et al*. 2016) during the submission process to the NCBI submission portal. Additional annotations, as well as annotations of *Thermanaeomonas* sp. strain 9S (DSM 22490) and the *T. burensis* MAG, were carried out with Prokka v1.14.6 (Seemann 2014) and with KofamKoala (Aramaki *et al*. 2020) against the KEGG database. Prediction of protein domains was performed with the module DomainAnnotation v.1.0.10 (against all domain libraries) as implemented in KBase (Arkin *et al*. 2018). Visualisation of gene clusters was done with R v4.3.2 in RStudio v2023.12.0+369 using the R package gggenes v0.5.1 (Wilkins 2020) as implemented in ggplot2 v3.4.4 (Wickham 2016).

### Phylogenetic analyses and taxonomic placement of strain AZ2

Taxonomic classification of strain AZ2 and assignment of lineage names were performed according to the Genome Taxonomy Database (GTDB) release R232 using GTDB-Tk v2.3.2 (Chaumeil *et al*. 2020, Parks *et al*. 2020). Genomes belonging to f_Neomoorellaceae (57 genomes) together with the genome of *Thermoanaerobacter kivui* (DSM 2030^T^) used as the outgroup, were obtained from GTDB R232. A set of 120 single-copy gene markers were identified and aligned with GTDB-Tk v2.4.0. The resulting alignment was trimmed with trimAl v1.4.1 using the -gappyout option (Capella-Gutiérrez, Silla-Martínez and Gabaldón 2009), yielding a final alignment of 4,794 bp. The best-fit-substitution model was determined with ModelFinder implemented in IQ-TREE v2.0.6 (Minh *et al*. 2020). A maximum-likelihood phylogenetic tree was inferred in IQ-TREE using the LG+F+R4 model with 1000 standard non-parametric bootstrap replicates. The tree was visualised in iTOL v7 (Letunic and Bork 2021).

Digital DNA-DNA hybridization (dDDH) values (and confidence intervals) between strain AZ2 and its closest relatives were calculated using the Type (Strain) Genome Server (https://tygs.dsmz.de) and the recommended settings of GGDC v4.0 (Meier-Kolthoff *et al*. 2013, 2022). Furthermore, average nucleotide identity (ANI) values were calculated using OrthoANI in the web service by CJ Bioscience (Yoon *et al*. 2017). Average amino acid identity (AAI) values were calculated with CompareM v0.1.2 (https://github.com/dparks1134/CompareM).

### Distribution of CODH gene clusters across members of the GTDB DSM-521 class

All genomes assigned to the GTDB class: c DSM-521 (65 genomes) were obtained from GTDB R232 (75 genomes) (Parks *et al*. 2020). A phylogenomic tree was reconstructed as described above, using 120 bacterial single-copy bacterial genes, a trimmed alignment of 4,895 bp, the LG+F+R4 substitution model and 1000x bootstrap replicates. To identify the genes encoding for CODHs, the genomes were re-annotated with Prokka v1.14.6 and candidate genes were manually confirmed with InterProScan v5.66-98 (Jones *et al*. 2014).

## Results and Discussion

### Enrichment and isolation of a novel thermophilic CO-utilising acetogen, strain AZ2

In this study, a thermophilic enrichment culture was developed from a marine sediment sample collected at a geothermal site in Ponta de Ferraria, São Miguel island, Azores, Portugal. The culture was grown under a gas phase of H_2_/CO_2_/CO (48:12:40% v/v, 1.7 atm). Following the initial 35-day incubation of the sediment sample, the culture completely consumed CO (23.5 mM) and produced acetate (8.9 mM) as the main end-product. Two subsequent 10% (v/v) dilutions enriched the culture toward a dominant Amplicon Sequence Variant (ASV) affiliated with *Thermanaeromonas sp.*, with a relative abundance of 91.3% (based on 16S rRNA amplicon sequencing) (Supplementary Figure S1). Strain AZ2 was isolated from this thermophilic enrichment after five subsequent 1% (v/v) serial dilutions (Supplementary Figure S2).

Strain AZ2 is an obligate anaerobic, thermophilic bacterium. It grew within a temperature range of 45 and 70°C, with an optimum at 65°C and a pH range of 5.5- 9.0 with an optimum at pH 7.0. It could grow fermentatively with D-glucose, D-galactose, D-fructose, pyruvate, formate, fumarate, succinate, producing, in most cases, acetate. When grown on succinate the fermentation-end product was propionate. Additionally, strain AZ2 could reduce thiosulfate and perchlorate with CO as an electron donor. Reduction of the electron acceptors was accompanied by acetate production. Acetate could not serve as an electron donor for the reduction of thiosulfate or perchlorate.

Furthermore, strain AZ2 could utilise and grow on CO as its sole energy and carbon source. When growing on N_2_/CO_2_/CO (48:12:40% v/v, 1.7 atm) (Figure 1a), or only CO (100% v/v, 1.7 atm) (Figure 1b), consumption of CO was accompanied by acetate production (7 mM and 15 mM, respectively). Trace amounts of H_2_ and propionate were also detected during growth of strain AZ2 on 100% CO. Propionate production from CO fermentation by a single strain has not been previously reported, and while only trace amounts were detected, its production was consistently observed across experiments.

**Figure 1.**
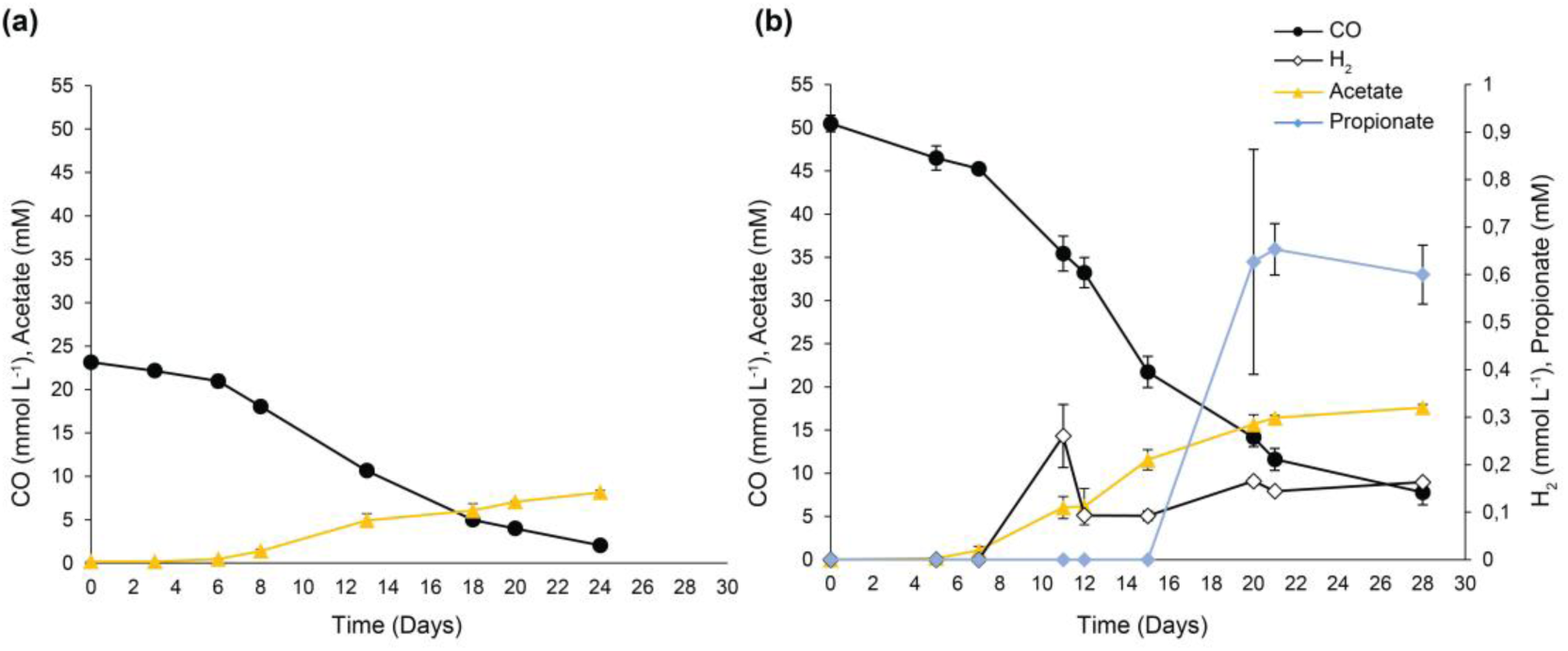
CO consumption and metabolite production by strain AZ2 under a gas phase of (a) N_2_/CO_2_/CO (48:12:40% v/v, 1.7 atm), and (b) CO (100% v/v, 1.7 atm) at 60°C. Error bars represent standard deviations of triplicate incubations.

### Phylogenetic position of strain AZ2 and reclassification of *Thermanaeromonas* sp. strain 9S (DSM 22490) and *Thermanaeromonas burensis* (DSM 26576^T^)

According to 16S rRNA gene analysis, strain AZ2 clusters within the *Thermanaeromonas* genus, which currently comprises two validly published species, *Thermanaeromonas toyohensis* (DSM 14490^T^) (Mori *et al*. 2002) and *T. burensis* (DSM 26576^T^) (Gam *et al*. 2016) and one uncharacterised species *Thermanaeromonas* sp. strain 9S (DSM 22490). The 16S rRNA gene of strain AZ2 shared a 93.5% sequence identity with the 16S rRNA gene of *T. toyohensis* DSM 14490^T^. This value falls within the 92-95% range, used for family delineation, but below the 95-98.6% range for genus delineation (Konstantinidis, Rosselló-Móra and Amann 2017), indicating that strain AZ2 likely belongs to a different genus than *T. toyohensis*.

Genome-based analysis confirmed the results of the 16S rRNA gene analysis. According to GTDB-Tk (GTDB R232), strain AZ2 was classified within the Neomoorellaceae family and the uncharacterised GTDB genus UBA 12545. To further investigate its taxonomic placement, we employed a phylogenomic approach using publicly available representative genomes and Metagenome Assembled Genomes (MAGs) belonging to the same family. We also sequenced the genomes of the other *Thermanaeromonas* isolates deposited in culture collections, *Thermanaeromonas* sp. strain 9S (DSM 22490) and *T. burensis* (DSM 26576^T^) and included them in the phylogenomic analysis. The genome of *T. toyohensis* (DSM 14490^T^) was already publicly available. Inference of a maximum-likelihood tree (created from a concatenated alignment of 120 single-copy bacterial phylogenetic markers) supported the placement of strain AZ2 in the UBA 12545 genus and not in the *Thermanaeromonas* genus, with high bootstrap values (99.9) (Figure 2).

**Figure 2.**
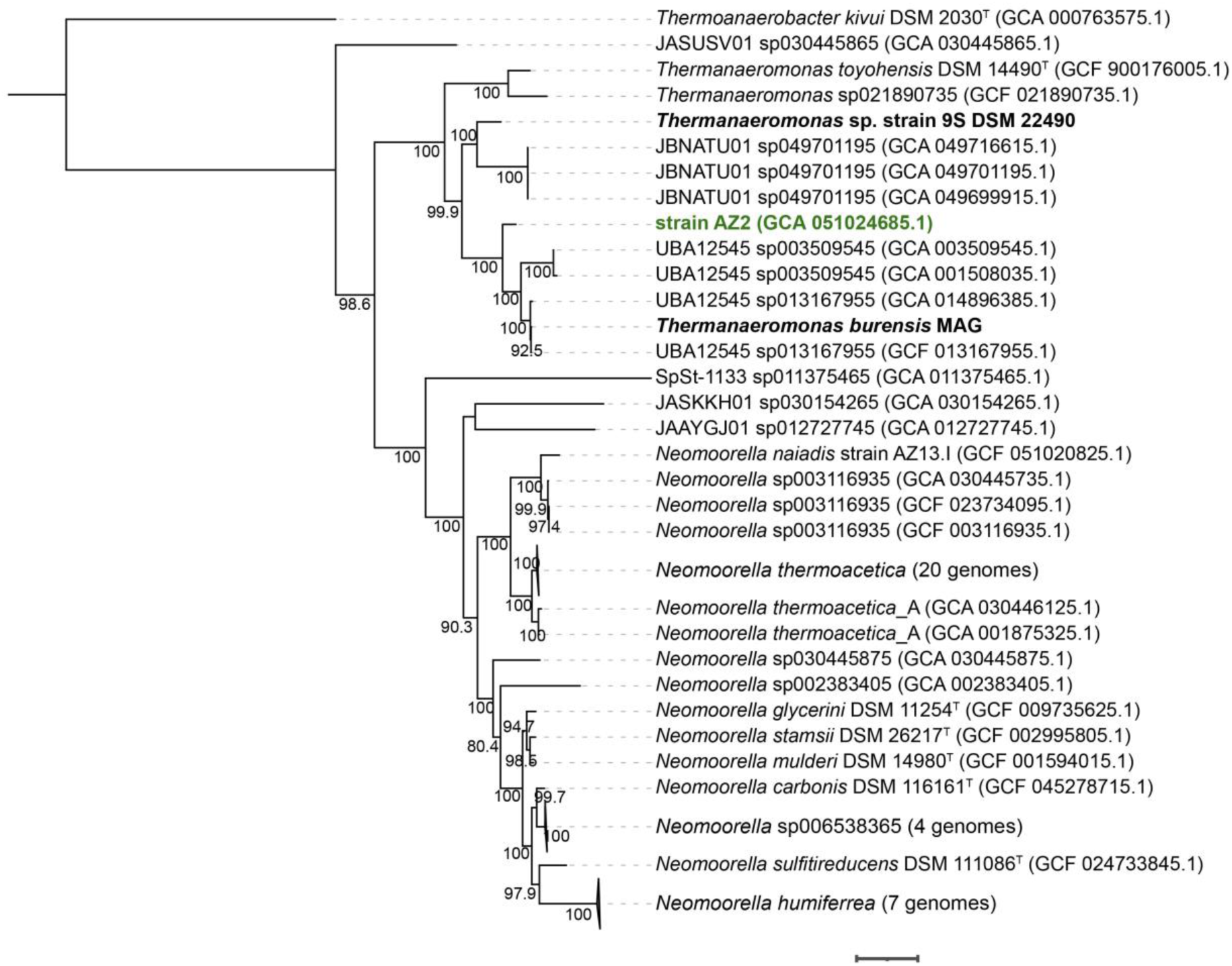
Maximum-likelihood phylogenetic tree based on the concatenated sequences of 120 single-copy bacterial phylogenetic marker genes. The scale bar represents 0.1 substitutions per site. The numbers at the nodes represent bootstrap values, and accession numbers are given in parentheses. Strain AZ2 is highlighted in green. Species names corresponding to genomes sequenced in this study are shown in bold. *Thermoanaerobacter kivui* was used as an outgroup and root of the tree.

The genome of *Thermanaeromonas* sp. strain 9S (DSM 22490) also did not cluster within the *Thermanaeromonas* genus, but instead within the JBNATU01 genus of the Neomoorellaceae family. While analysing the genome of *T. burensis* (DSM 26576^T^), we detected contamination of the DSMZ-provided culture. After binning the contigs, we recovered two MAGs from the DNA sample (Supplementary Table S1). One of the MAGs represented *T. burensis,* and we will refer to it as *T. burensis* MAG from now on. The second MAG represented an uncultured *Geochorda* sp. The *T. burensis* MAG was used for further genomic analyses, and similarly to strain AZ2 it was placed in the UBA 12545 genus.

Furthermore, the dDDH and ANI values between strain AZ2 and the rest of the UBA 12545 members, including *Thermanaeromonas* sp. strain 9S (DSM 22490) and the *T. burensis* MAG were between 23.4-32.5% and 78.4-86.7%, respectively (Supplementary Table S2). Both value ranges fulfil the criteria for prokaryotic species delineation, proposed to be >70% (dDDH) and >95-96% (ANI) (Konstantinidis et al., 2017; Chun et al., 2018; Jain et al., 2018).

Therefore, based on the phenotypic (Supplementary Table S3), genotypic and phylogenetic characteristics, we conclude that strain AZ2 represents a new species in the uncharacterised UBA 12545 genus. We propose the name *Thermobium azorense* gen. nov., sp. nov for isolate AZ2.

### Genome properties of strain AZ2

The draft genome of strain AZ2 was assembled into 92 contigs. The genome size is 2,970,900 bp (N50 value: 52,920 bp), and the in silico-determined G+C content of the genome is 55.14 mol%. The genome is 99.49% complete and has low contamination (0.58%). A total of 2,913 genes were predicted in the genome, including 2,837 protein-coding genes, 53 RNA genes (3 rRNAs, 46 tRNAs, and 4 ncRNAs) and 23 pseudogenes.

The genome of strain AZ2 harbours all the genes required for the WLP (Figure 3), typically found in acetogens for carbon fixation from either CO_2_ or CO. While strain AZ2 could indeed grow with CO as its sole carbon and energy source, it failed to grow on CO_2_/H_2_ even after two months of incubation. The inability of other *Neomoorella* spp., which belong to the same family as strain AZ2, to grow on CO_2_/H_2_ has been previously reported (Slobodkin *et al*. 1997; Nepomnyashchaya *et al*. 2012; Alves *et al*. 2013; Slobodkina *et al*. 2022; Galani *et al*. 2025). All genes required for glycolysis were found in the genome of strain AZ2. Although most genes for the TCA cycle were present, genes encoding fumarate reductase, malate dehydrogenase or succinyl-CoA synthetase were not found. Strain AZ2 can produce propionate under laboratory conditions when grown on succinate, potentially utilising the succinate pathway. The genome includes all the genes required for this pathway, except for the gene encoding succinyl-CoA synthetase (catalysing the interconversion between succinate and succinyl-CoA). This gene may have been misannotated or located in a misassembled region of the genome. Alternatively, strain AZ2 might employ a CoA-transferase to perform the interconversion of succinate to succinyl-CoA. The genome contains two genes predicted to encode CoA transferases (V3F56_05410 and V3F56_05370). Specifically, the latter one shares a 91.75% identity with succinyl-CoA:acetate CoA- transferase of *Thermanaeromonas* sp. C210. This enzyme could catalyse the reversible transfer of CoA between acetyl-CoA and succinate, resulting in the formation of succinyl-CoA and acetate, providing a potential CoA-transferase-mediated route for the conversion of succinate to succinyl-CoA (Pettinato, Böhnert and Berg 2022).

**Figure 3.**
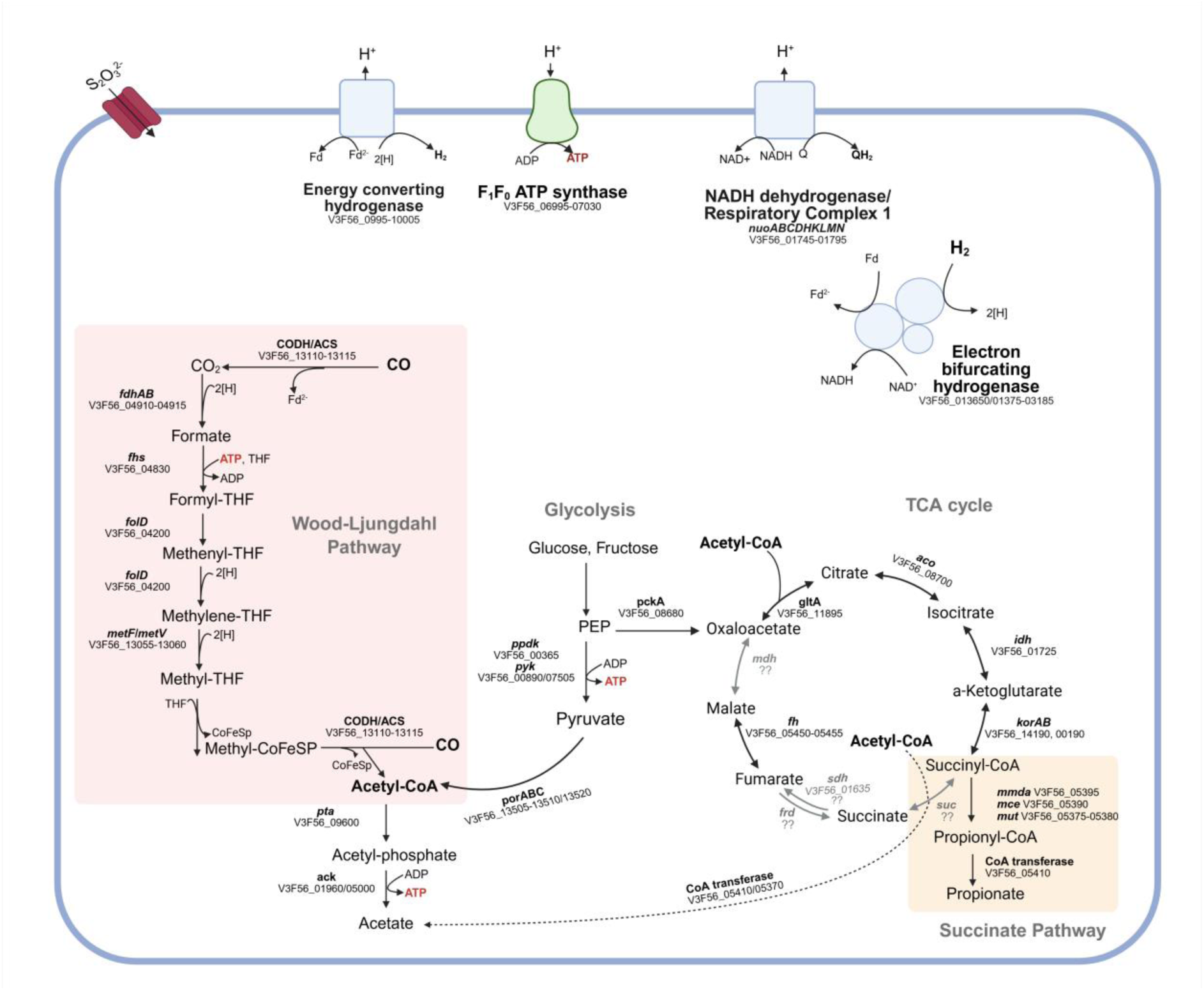
Key genes and metabolic pathways identified in the genome of strain AZ2. Gene abbreviations: CODH-ACS: CO-dehydrogenase acetyl-CoA synthase; fdhAB: formate dehydrogenase; fhs: formate--tetrahydrofolate ligase; folD: methylene tetrahydrofolate dehydrogenase; metF-metV: methylenetetrahydrofolate reductase; pta: phosphotransacetylase; ack: acetate kinase; porBC: pyruvate:ferredoxin oxidoreductase; ppdk: pyruvate phosphate dikinase; pyk: pyruvate kinase; pckA: PEP carboxykinase; gltA: citrate synthase; aco: aconitase; idh: isocitrate dehydrogenase; korAB: α-ketoglutarate oxidoreductase; mmdA: methylmalonyl-CoA decarboxylase; mce : methylmalonyl-CoA epimerase; mut: methylmalonyl-CoA mutase; CoA trasferase: acetyl-CoA transferase; suc: succinyl-CoA synthetase; sdh: succinate dehydrogenase; frd: fumarate reductase; fh: fumarate hydrogenase; nuo: NADH:quinone oxidoreductase (Respiratory Complex 1). Figure created with *BioRender*.

Two potential energy conservation systems were found in strain AZ2, an energy-converting hydrogenase (Ech) and a NADH dehydrogenase-like enzyme that is closely related to Respiratory complex 1 (encoded by *NuoABCDHKLMN*). While Ech is one of the major energy conservation mechanisms found in acetogens, the NADH dehydrogenase-like complex has only recently been proposed to contribute to energy conservation in *Neomoorella* species and remains poorly characterised (Santaella, Sousa and Stams 2025; Rosenbaum *et al*. 2026). Similar to the corresponding complexes in *N. thermoacetica* and *N. caeni*, the complex found in the genome of strain AZ2 lacks the NADH binding module (*NuoEFG*). In these species, the complex has been hypothesised to employ ferredoxin rather than NADH as electron donor, coupling menaquinone reduction to ion translocation across the membrane. The presence of a highly similar complex in strain AZ2 suggests that it may perform a similar physiological role. Therefore, both systems are likely involved in proton translocation across the membrane, which fuels the H^+^-dependent F_1_F_0_ ATP synthase (Rosenbaum and Müller 2021, 2023). Additionally, the genome of strain AZ2 encodes a putative electron bifurcating hydrogenase (HydABC), which couples H_2_ oxidation with the reduction of NAD^+^ and ferredoxin, subsequently used in the WLP (Huang *et al*. 2012; Wang *et al*. 2013).

### *Thermanaeromonas* species are capable of CO conversion

To date, *Thermanaeromonas* species have not been systematically studied for their ability to utilise CO, and thus, this metabolic capability has been overlooked. A draft genome of *Thermanaeromonas* sp. C210 (Inoue *et al*. 2020), reported to have been isolated on CO, has been published. However, no corresponding strain is available in culture collections, and the genome announcement remains the only available record. In this study, we tested the ability of *T. toyohensis* (DSM 14490^T^) (Mori *et al*. 2002) and *Thermanaeromonas* sp. strain 9S (DSM 22490) to use CO as a growth substrate. Our findings revealed that both of them could convert CO; nevertheless, their production profiles from CO fermentation differed from the one observed in strain AZ2. Both *T. toyohensis* (DSM 14490^T^) and *Thermanaeromonas* sp. strain 9S (DSM 22490) produced mainly H_2_ when growing on CO (Figure 4). *T. toyohensis* (DSM 14490^T^) also produced trace amounts of acetate (0.9 mM). The culture of *T. burensis* (DSM 26576^T^) (Gam *et al*. 2016) (contaminated with a *Geochorda* bacterium) was also tested and could grow on CO, producing acetate as the main end-product (Supplementary Figure S3).

**Figure 4.**
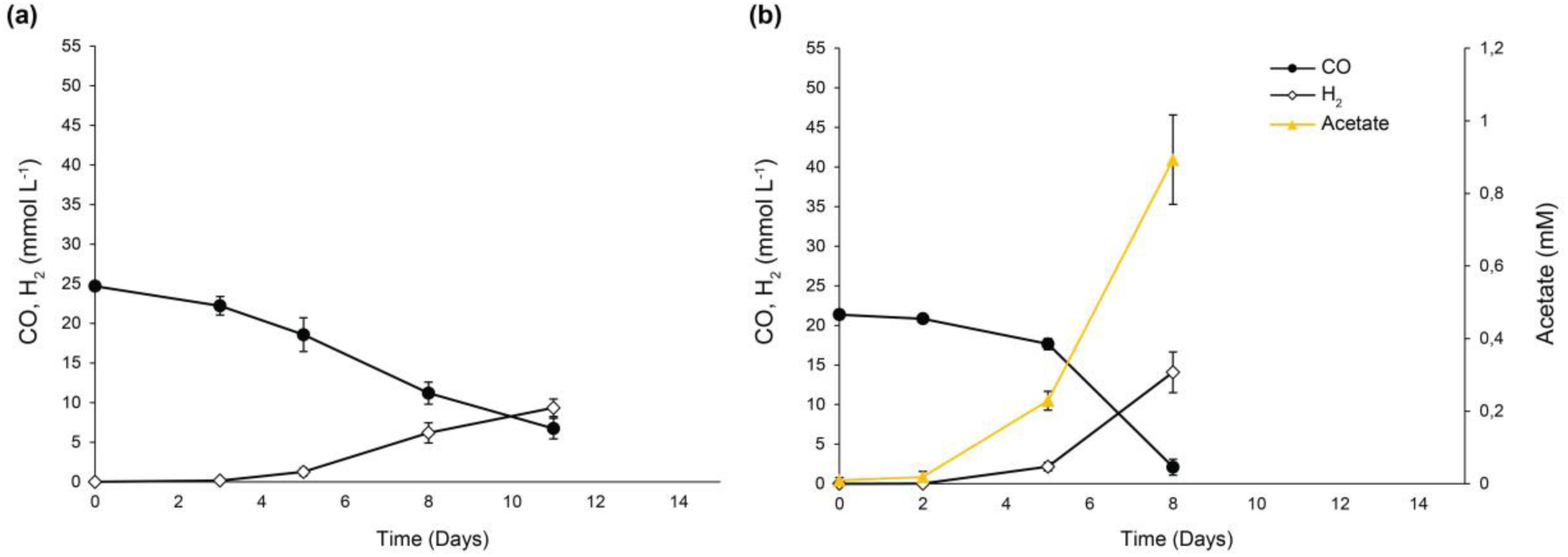
CO consumption and metabolite production by (a) *T. toyohensis* (DSM 14490^T^), (b) *Thermanaeromonas* sp. strain 9S (DSM 22490). Incubations were done under an N_2_/CO_2_/CO (48:12:40% v/v, 1.7 atm) headspace, at 70°C and 60°C, respectively. Error bars represent standard deviations of triplicate incubations.

### Comparative genomic analyses of *cooS* genes encoding for CODHs in Thermanaeromonas *spp*

Genomic comparison of strain AZ2 with other *Thermanaeromonas* species showed that all genomes featured *cooS* genes, which encode the catalytic domain of anaerobic CODHs, but in different copy numbers and located in distinct gene clusters (Figure 5).

**Figure 5.**
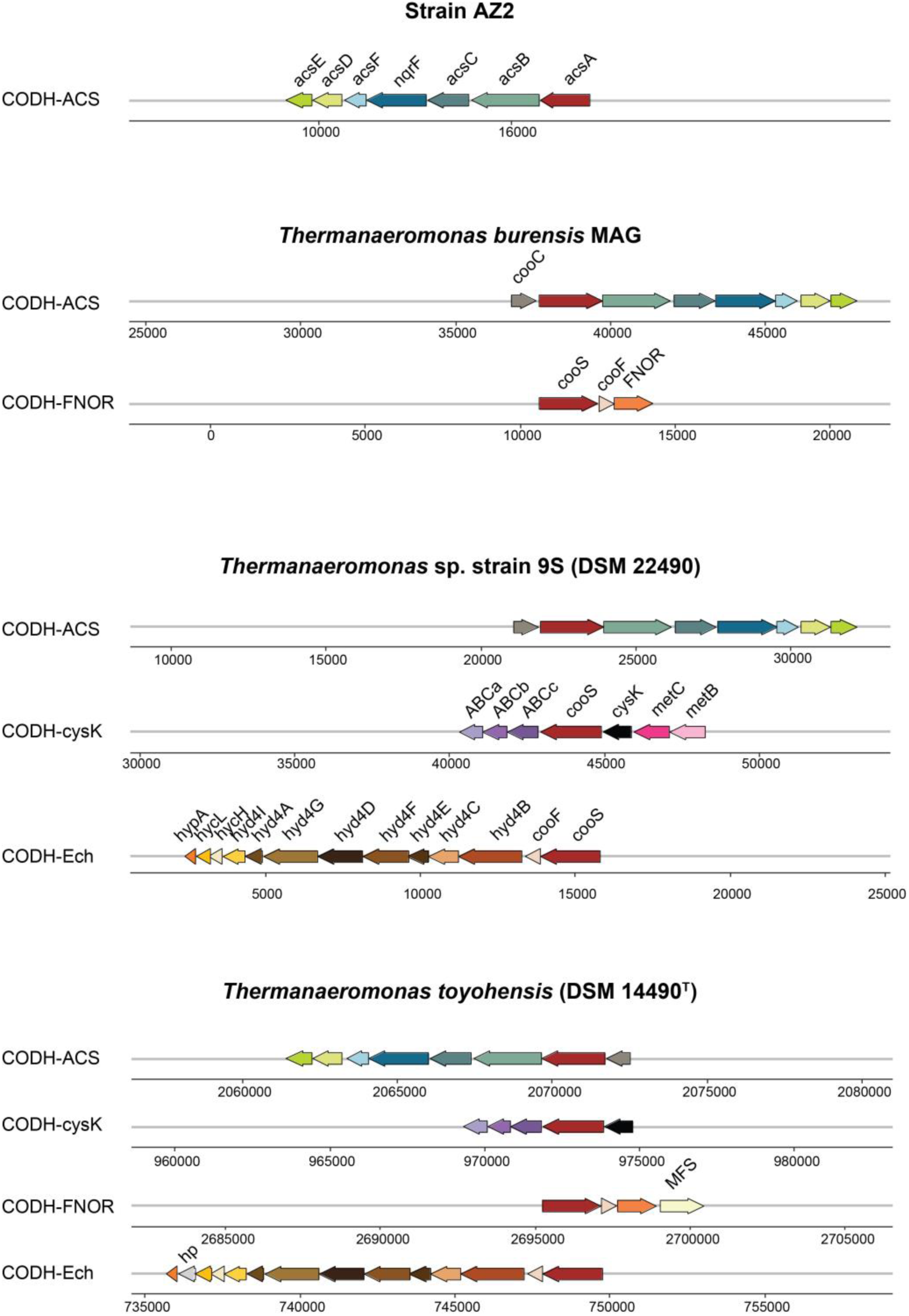
Genomic structure of CODH gene clusters in strain AZ2 and its closest isolated relatives, *T. burensis* MAG, *Thermanaeromonas* sp. strain 9S (DSM 22490) and *T. toyohensis* (DSM 14490^T^). Gene abbreviations: cooC: CODH chaperone; acsA or cooS: CO-dehydrogenase; acsB: Acetyl-CoA synthase; acsC: corrinoid iron–sulfur protein large subunit; acsD: corrinoid iron–sulfur protein small subunit; acsE: methyltransferase A; acsF: ACS chaperone; nqrF: Na(+)-translocating NADH-quinone reductase subunit F, cooF: ferredoxin-like electron transfer Fe-S protein, FNOR: NAD/FAD oxidoreductase, ABCa: ABC transporter domain a, ABCb: ABC transporter domain b, ABCc: ABC transporter domain c, cysK: cysteine synthase, metC: cystathionine beta-lyase, metB: cystathionine gamma-lyase, hycL: hydrogenase maturation peptidase, hycH: hydrogenase maturation protein,hyd4A-I: NiFe hydrogenase group 4 subunits A-I, MFS: major facilitator superfamily transporter, hp: hypothetical protein.

The genome of strain AZ2 contains a single copy of the *cooS* gene (*acsA*, V3F56_13115) located within a CODH-ACS cluster. However, the *cooS* gene appears to be fragmented as it is found at the end of a contig. We assume that its shorter size (330 bp) results from assembly issues. The CODH-ACS gene cluster encodes the bifunctional CODH-ACS, which is responsible for acetyl-CoA generation, at the last step of the WLP. Acetyl-CoA can further be converted into acetate, explaining the observed acetate production when strain AZ2 is growing on CO.

*T. burensis* MAG contained two *cooS* gene copies: one in a cluster with ACS, and the other adjacent to genes encoding a ferredoxin-like electron transfer protein (CooF) gene and a FAD/NAD(P) oxidoreductase (FNOR). The *T. toyohensis* genome contained a similar CODH-FNOR gene cluster. Several functions have been proposed for this gene cluster. In *C. hydrogenoformans*, where it was first discovered, as well as in mesophilic Clostridia (e.g. *Clostridium autoethanogenum* and *C. ljungdahlii)*, this gene cluster also includes a gene coding for a rubrerythrin-like protein leading to the hypothesis of its involvement in response to oxidative stress (Wu *et al*. 2005; Hillmann *et al*. 2009; Whitham *et al*. 2015; Liew *et al*. 2016a). Both genomes of *T. burensis* MAG and *T. toyohensis* lack the rubrerythrin coding gene. In *Geobacter* spp., which also lack this gene, the CODH-FNOR is predicted to be involved in energy conservation via CO oxidation coupled to NAD(P)^+^ reduction (Geelhoed, Henstra and Stams 2016).

The genomes of *Thermanaeromonas* sp. strain 9S (DSM 22490) and *T. toyohensis* (DSM 14490^T^) both had one *cooS* gene copy in a cluster with ACS as well as one copy in a cluster with genes encoding domains of an energy-converting hydrogenase (Ech). This CODH-Ech cluster is found in CO-oxidising anaerobes capable of coupling CO oxidation to proton reduction, leading to the production of H_2_ and CO_2_ (Sokolova *et al*. 2009; Sant’Anna *et al*. 2015; Omae *et al*. 2017; Fukuyama *et al*. 2020). This could explain the observed H_2_ production by these bacteria during growth on CO. Furthermore, again in both genomes of *Thermanaeromonas* sp. strain 9S (DSM 22490) and *T. toyohensis*, there is one *cooS* copy next to a gene encoding for cysteine synthase (cysK) and genes encoding for the three domains of an ABC transporter. In the genome of *Thermanaeromonas* sp. strain 9S (DSM 22490) next to the CODH-cysK gene cluster, there were also two genes encoding for domains of cystathionine lyase, while these genes were absent from the genome of *T. toyohensis* (DSM 14490^T^). A similar gene cluster has also been found in the genomes of *Neomoorella thermoacetica* and *Calderihabitans maritimus* (Omae *et al*. 2017), though its function remains elusive.

When comparing the production profiles with the number of *cooS* gene copies and their genomic context in the genomes of *Thermanaeromonas* spp., a pattern emerges. In bacteria like strain AZ2, which possess only the CODH-ACS cluster, acetate is the primary product of CO utilisation. However, when the CODH-ACS cluster coexists with the CODH-Ech cluster, the production profile shifts towards H_2_ production. This shift is likely because the hydrogenogenic metabolism is significantly faster than acetogenesis from CO (Diender, Stams and Sousa 2015), providing the bacteria with a rapid mechanism to mitigate CO toxicity. The other two types of CODH-gene clusters (CODH-FNOR and CODH-cysK) have not been shown to lead to the production of specific compounds, and their metabolic role is still not fully understood.

### Novel carboxydotrophic potential within the GTDB class DSM-521

Following the comparative genomic analysis of the *cooS*-gene clusters in *Thermanaeromonas* spp., we expanded our investigation to include the entire Moorellia class, to which strain AZ2 belongs. The GTDB class DSM-521 currently includes three families (f_JACIYF01, f_SLTJ01, f_Neomoorellaceae) represented both by MAGs and isolates, retrieved from different types of hot (in most cases) environments (Figure 6). Most of the isolates belong to the f_Neomoorellaceae family, which includes the best-characterised thermophilic CO-utilising genus *Neomoorella* (Galani *et al*. 2025).

**Figure 6.**
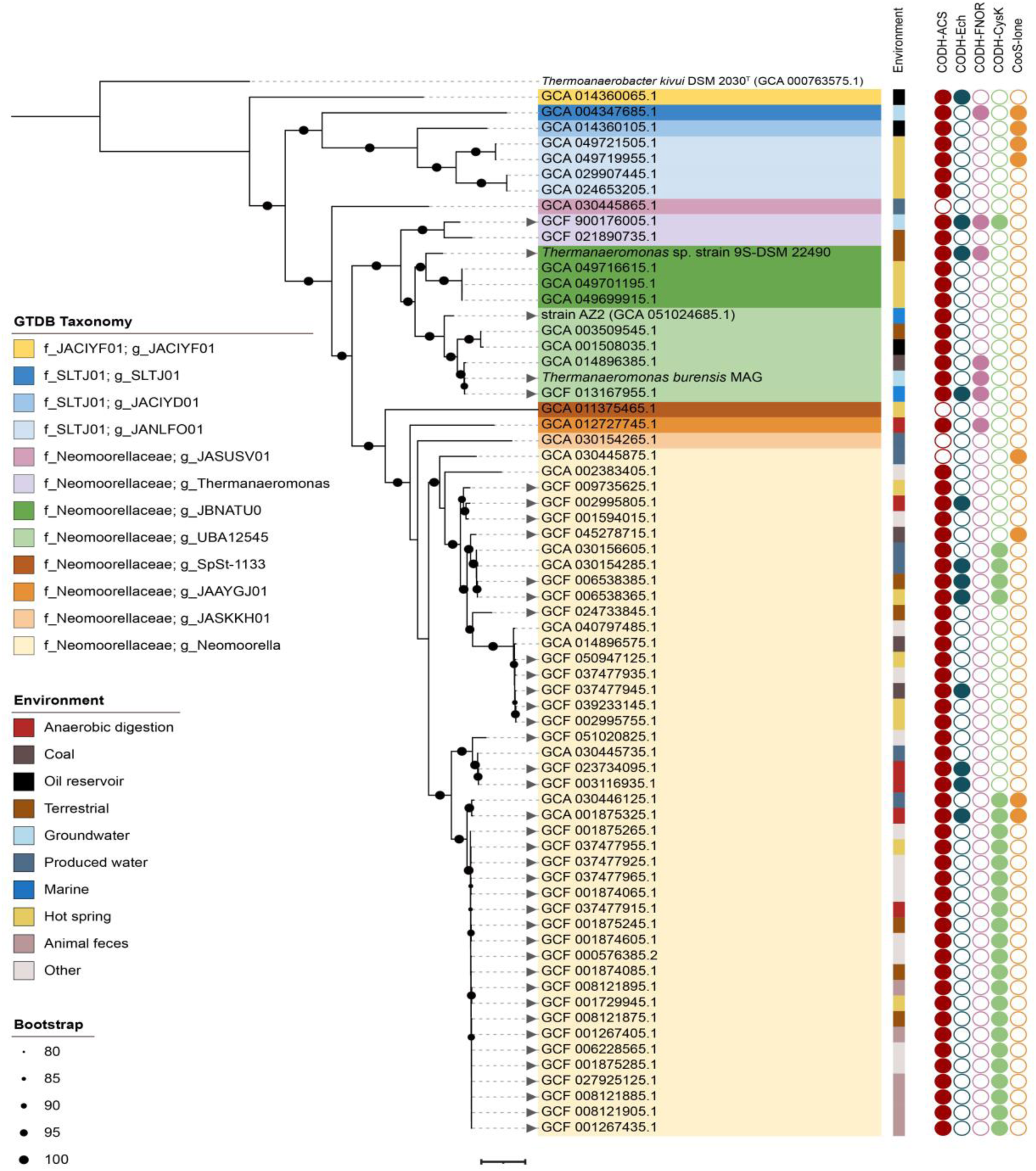
Phylogenomic tree of the GTDB DSM-521 class, based on the concatenated sequences of 120 single-copy bacterial phylogenetic marker genes. Bootstrap values over 80% are shown by black circles. Triangles before the accession numbers represent isolates. Different colours on the clades represent assigned taxonomic classification at the genus level, following the GTDB (R2323) taxonomy. Coloured strips next to the clades indicate the environmental isolation source. Coloured and non-coloured circles indicate the presence or absence of CODH gene clusters. The scale bar represents 0.1 substitutions per site. *Thermoanaerobacter kivui* was used as an outgroup and root of the tree.

Our findings suggest that the genomic potential for CO utilisation (as indicated by the presence of *cooS* genes) is widespread among the members of the DSM-521 class. Among the genomes analysed, only three completely lacked *cooS* genes, all three of them belonging to the f_Neomoorellaceae family (g_SpSt-1133, g_JASKKH01, g_JASUSV01 genera).

Furthermore, most of the genomes featured multiple *cooS* copies, suggesting that CO could be an important energy or carbon source in their respective habitats. In recent years, numerous species have been found to encode multiple CODHs. While not all of them have clearly defined functional roles, it is likely that the presence of multiple CODH gene clusters enhances the organism’s fitness in CO-rich environments by offering diverse metabolic pathways for CO utilisation (Techtmann *et al*. 2012). Additionally, it could provide a growth advantage to the organism, as the various CODHs may exhibit different affinities for distinct CO concentrations.

The most frequently identified gene cluster in the DSM-521 class was the CODH-ACS, suggesting that most microbes in this class may have the ability to utilise CO as a carbon source through the WLP. The CODH-Ech gene cluster was predominantly found in previously isolated microbes known for primarily producing H_2_ from CO oxidation, such as *Neomoorella caeni* and *N. stamsii* (Alves *et al*. 2013; Santaella, Sousa and Stams 2023), further confirming the pattern observed in the *Thermanaeromonas* spp. The CODH-FNOR and CODH-CysK gene clusters were also found in a lot of genomes, but in most cases, they co-existed with the CODH-ACS or Ech ones, acting as secondary CODHs.

At the same time, several genomes analysed in this study featured *cooS* genes (“lone *cooS’’*) that were not part of already characterised gene clusters and no assumption for their function can be ascribed based on the genes that are in their proximity. Most of the lone *cooS* genes were found in MAGs and further studies are needed to elucidate their role.

Most genomes in the DSM-521 class carry the CODH-ACS gene cluster without, in most cases, co-occurrence of the CODH-Ech gene cluster. Combined with the observed patterns linking these clusters to the production of specific products (as discussed in the previous section), this suggests that several non-isolated members of this class could represent novel carboxydotrophic acetogens. These microbes are promising targets for future isolation and characterisation approaches. Utilising genomic data to predict the functions of CODHs and the production profiles of microbes can significantly aid in identifying novel microorganisms capable of utilising CO and exhibiting the desired production profiles for biotechnological applications. This is particularly important for thermophilic acetogens because, despite their numerous advantages for industrial syngas fermentation, their application remains limited due to the small number of isolates currently available.

### Description of *Thermobium* gen. nov

(Ther.mo’bi.um. Gr. masc. adj. thermos, hot: Gr. masc .n. bios, life; N.L. neut. n. Thermobium, life form from a hot environment). Cells are rod-shaped and stain Gram-positive. Spore-forming. Strict anaerobes and moderately thermophilic. They can perform fermentative acetogenic growth on sugars or on CO. Acetate is the main end product from fermentation. The type species is *Thermobium azorense*.

### Description of *Thermobium azorense* gen. non, sp. nov

*Thermobium azorense* (a.zo.ren’se. N.L. neut. adj. azorense, pertaining to the Azores).

Cells are straight rods, 0.5-0.6 μm wide and 2-3 μm long, and occur single or in pairs. Cells stain Gram-positive. After prolonged times of incubation at 60°C, cells produce terminal round endospores. It is an obligate anaerobic, thermophilic bacterium. Growth occurs between 45 and 70°C with an optimum at 65°C (at pH 7.0). The pH range for growth is 5.5-9.0 with an optimum pH of 7.0. Growth does not occur above a NaCl concentration of 2.5% (w/v). Yeast extract was not required for growth.

It grows fermentatively with D-glucose, D-galactose, D-fructose, pyruvate, formate, succinate, and CO [up to 100% CO (v/v), 1.7 atm]. In most cases the fermentation end-product is acetate. During the fermentation of succinate, propionate is the end product. Does not utilise xylose, maltose, sucrose, mannose, lactose, butyrate, ethanol, methanol, lactate, and H_2_/CO_2._ In the presence of CO as an electron donor, it reduces thiosulfate and perchlorate, with acetate as a by-product. Sulfate, fumarate and nitrate are not utilised as electron acceptors. Acetate does not serve as an electron donor for the reduction of thiosulfate or perchlorate.

The genome size of strain AZ2 is 2,970,900 bp (N50 value: 52,920 bp) with an in silico-determined G+C content of 55.14 mol%.

The type strain, AZ2^T^ (= DSM 121889^T^ = JCM 39698^T^), was isolated from marine sediment collected at Ponta de Ferraria on São Miguel island, Azores, Portugal.

## Conclusions

In the present study, we isolated a novel thermophilic CO-utilising acetogen, strain AZ2, which we propose as the first representative species of a novel genus, *Thermobium azorense* gen. nov sp. nov. Isolate AZ2 was capable of utilising CO, producing acetate as the main metabolic product. When grown on 100% CO, strain AZ2 produced trace amounts of H_2_ and propionate. We further showed that the closest isolated relatives of strain AZ, *T. toyohensis* and *Thermanaeromonas* sp. strain 9S, can also utilise CO, producing H_2_ as the end-product. This variation in production profiles could be linked to differences in their genomes where distinct *cooS*-gene clusters were identified. Strain AZ2 featured a CODH-ACS gene cluster, while *Thermanaeromonas* species featured both CODH-ACS and CODH-Ech gene clusters. This observation highlights the value of using genomic data to predict the production profiles of novel CO-utilising microbes. Finally, we explored the presence and organisation of *cooS*-gene clusters in all members of the DSM-521 class. Our findings revealed that the vast majority of microbes in this class have the genomic potential to utilise CO, likely linked to acetate production based on the widespread presence of the CODH-ACS gene cluster. Overall, this study provides insights for the targeted isolation of novel CO-utilising thermophiles. At the same time, the isolation of strain AZ2 expands the currently limited number of thermophilic acetogens, contributing to their further development and application in syngas fermentation.

## Funding

This study is part of Perspectief Programma P16-10 with project number 1 of the research programme MicroSynC, which is (partly) financed by the Dutch Research Council (NWO-TTW). Sampling for this study was supported by a KNAW Ecology fund (KNAWWF/747/ECO2021-8).

## Conflict of interest

The authors declare no conflict of interest.

## Data Availability

The Whole Genome Shotgun project for strain AZ2 has been deposited at DDBJ/ENA/GenBank under the accession JAZHPA000000000 (BioSample: SAMN39671199). The version described in this paper is version JAZHPA010000000. The GenBank accession number is GCA_051024685.

The Whole Genome Shotgun project for *Thermanaermonas* sp. (DSM 22490) has been deposited at DDBJ/ENA/GenBank under the accession number: JBJHHE000000000 (BioSample: SAMN44672059). The GenBank accession number is GCF_056378755.1. Both genomes were deposited under the BioProject PRJNA1037001.

The full 16S rRNA gene sequence of strain AZ2 was deposited in GenBank under the accession number PP598622.

## Notes

### Competing Interest Statement

The authors have declared no competing interest.

