## Supplementary files for "Carbon monoxide utilisation by *Thermanaeromonas* species and description of *Thermobium azorense* gen. nov., sp. nov."

**Supplementary materials**

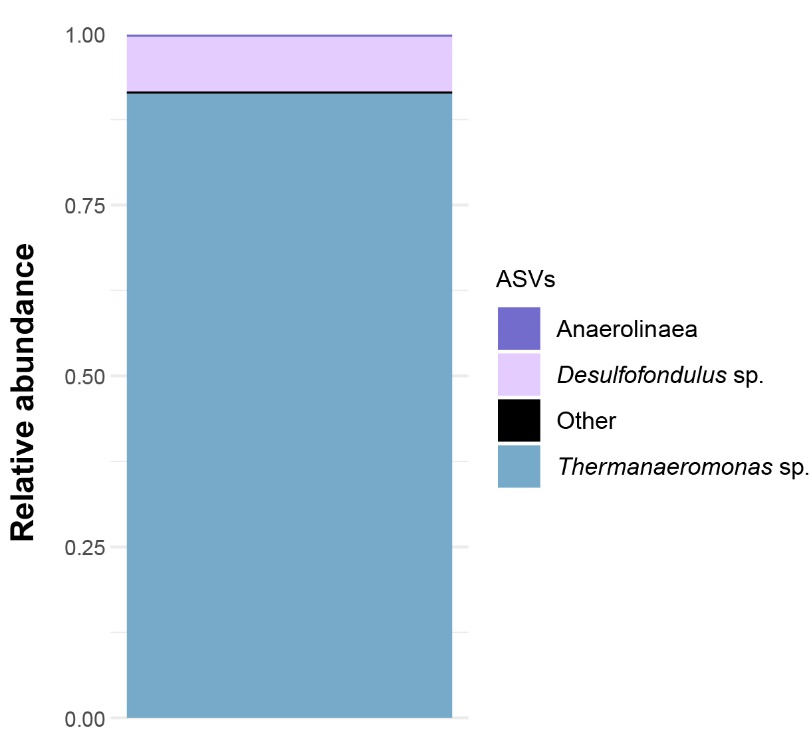

Supplementary Figure S1. 16S rRNA amplicon sequencing results of enrichment AZ2.

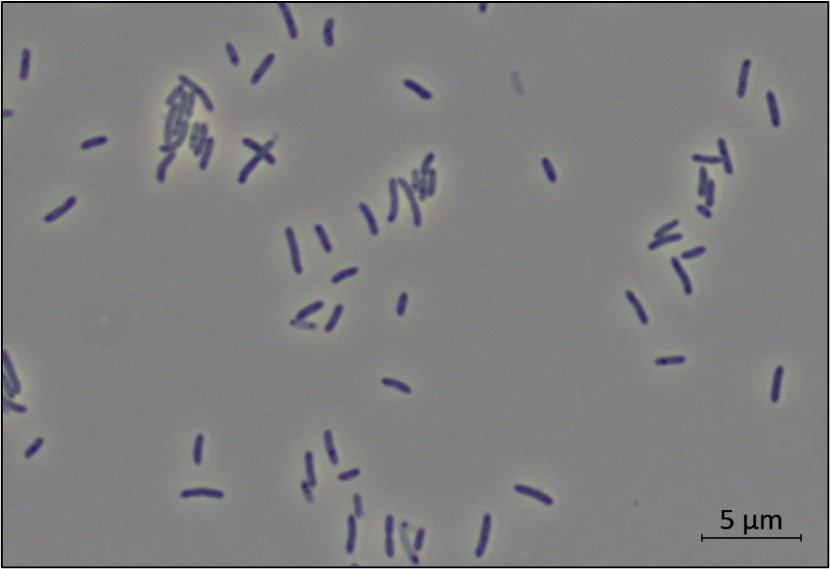

Supplementary Figure S2. Morphology of strain AZ2 grown on CO as observed by light microscopy. Scale bar = 5 μm.

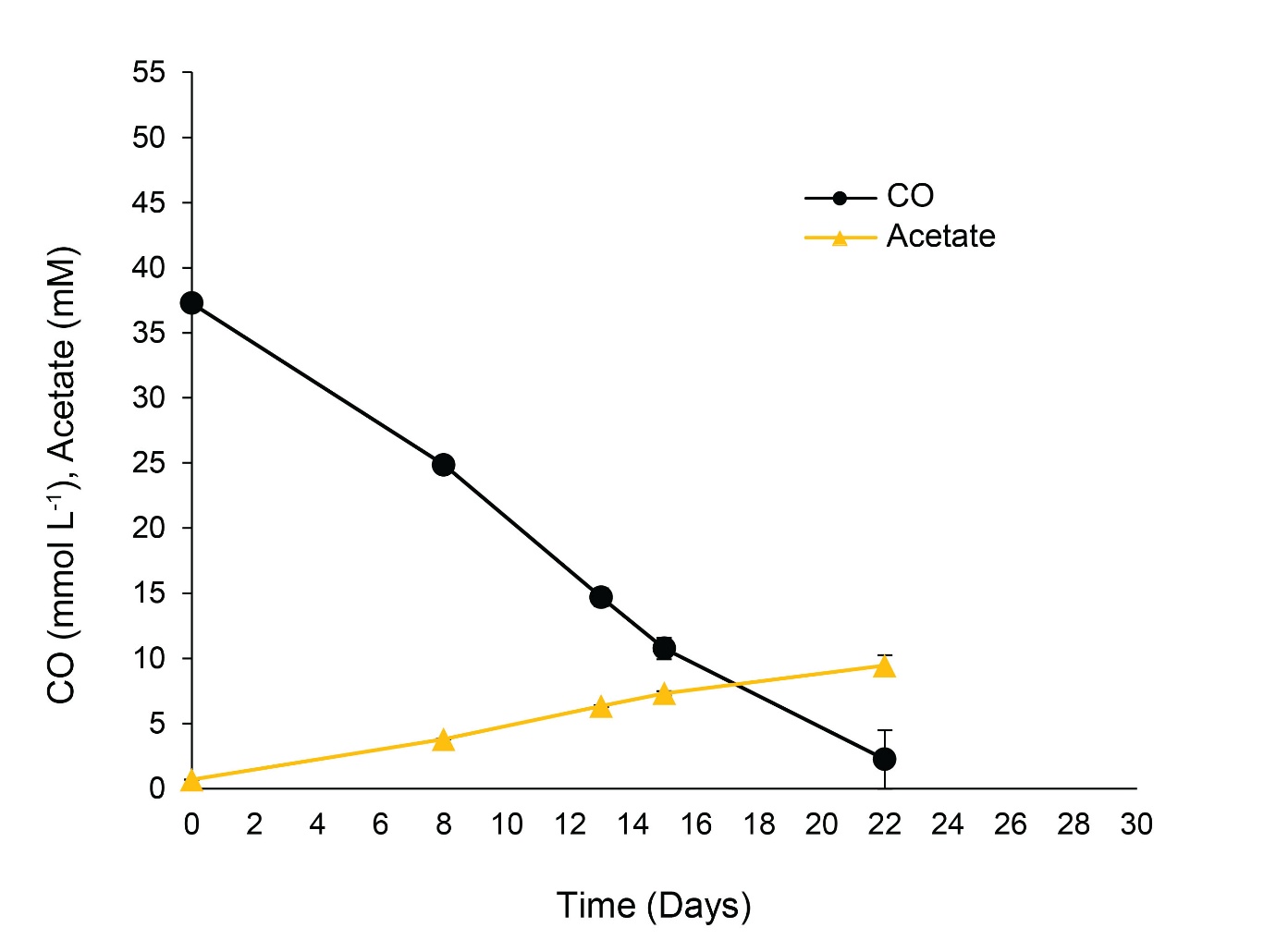

Supplementary Figure S3. CO consumption and metabolite production by T. burensis (DSM 26576T) under a gas phase of N2/CO2/CO (48:12:40% v/v, 1.7 atm) at 65oC. Error bars represent standard deviations of duplicate incubations.

Supplementary Table S1. Binning results of DSMZ culture-*Thermanaeromonas burensis* (DSM 26576^T^). Relative abundance values are expressed as the percentage of reads mapping to each bin relative to the total number of reads sequenced in the sample.

| **Bins** | **Completeness (%)** | **Contamination (%)** | **Relative abundance (%)** | **Taxonomy** |
| --- | --- | --- | --- | --- |
| Bin.1 | 100 | 0 | 19.38 | d__Bacteria; p__Bacillota; c__Limnochordia; o__Limnochordales; f__Geochordaceae; g__Geochorda; s__ |
| Bin.2 | 89.70 | 0.524 | 80.62% | d__Bacteria; p__Bacillota; c__DSM-521; o__Neomoorellales; f__Neomoorellaceae; g__UBA12545; s__UBA12545 sp013167955 |

Supplementary Table S2. Pairwise ANI, dDDH and AAI comparisons between strain AZ2 (GCA_051024685) and genomes of type strains belonging to the GTDB UBA12545 and *Thermanaeromonas* genera. C.I values refer to confidence intervals. * represent the GTDB representative species.

| **Organism** | **GenBank accession number** | **OrthoANIu (%)** | **dDDH (d4) [C.I.] (%)** | **Mean AAI (standard deviation) (%)** |
| --- | --- | --- | --- | --- |
| UBA12545_sp003509545* | GCA_001508035.1 | 82.57 | 26.6 [24.2 - 29.1] | 85.63 [10.05] |
| UBA12545_sp003509545 | GCA_001508035.1 | 81.86 | 25.2 [22.9 - 27.7] | 85.28 [9.68] |
| UBA12545_sp013167955* | GCA_013167955.1 | 86.75 | 32.5 [30.1 - 35.0] | 88.78 [9.59] |
| UBA12545_sp013167955 | GCA_014896385.1 | 86.42 | 31.6 [29.2 - 34.1] | 87.92 [10.62] |
| *Thermanaeromonas* sp. strain 9S (DSM 24990) | GCF_056378755.1 | 78.39 | 23.4 [21.1 - 25.9] | 81.10 [12.29] |
| *Thermanaeromonas burensis* MAG | - | 86.68 | 31.9 [29.5 - 34.4] | 88.56 [9.67] |
| Thermanaeromonas *toyohensis* (DSM 14490^T^) | GCA_900176005.1 | 71.99 | 24.3 [21.9 - 26.7] | 74.18 [12.8] |
| *Thermanaeromonas* sp021890735 | GCA_021890735.1 | 71.31 | 22.3 [20.0 - 24.8] | 73.76 [12.62] |

Supplementary Table S3: Distinctive characteristics of *Thermanaeromonas* strains. Characteristics of *T. toyohensis* (DSM 14490^T^) were obtained from Mori *et al*. (2002) and for and *T. burensis* (DSM 26576^T^) from Gam *et al*. (2016). +, positive; −, negative; n.r., not reported. * The ability of CO utilisation of *T. toyohensis* (DSM 14490^T^) and *T. burensis* (DSM 26576^T^) was confirmed in the present study. ^ *T. toyohensis* (DSM 14490^T^) cannot ferment formate, but it can oxidise it in the presence of thiosulfate as an electron acceptor. In the deposited culture of *T. burensis* (DSM 26576^T^) contamination was detected.

|  | **Strain AZ2** | ***T. toyohensis* (DSM 14490^T^)** | ***T. burensis* (DSM 26576^T^)** |
| --- | --- | --- | --- |
| **Isolation Source** | marine hydrothermal sediment | geothermal  aquifer | subterranean  clay formation |
| **Temperature ^o^C (optimum)** | 40-70 (65) | 55-73 (70) | 66-70 (65) |
| **pH (optimum)** | 5.5-9.0 (7.0) | 5.5-8.5 (6.5) | 5.5-9.0 (7.5) |
| **Electron donor/Carbon source** | | | |
| CO | + | +* | +* |
| H_2_/CO_2_ | - | - | - |
| Glycerol | - | n.r. | n.r. |
| Glucose | + | + | + |
| Galactose | + | - | n.r. |
| Fructose | + | + | - |
| Maltose | - | + | + |
| Sucrose | - | + | n.r. |
| Xylose | - | + | - |
| Mannose | - | + | - |
| Lactose | - | - | - |
| Pyruvate | + | + | n.r. |
| Acetate | - | - | - |
| Formate | + | + ^ | - |
| Propionate | - | n.r. | n.r. |
| Butyrate | - | n.r. | n.r. |
| Lactate | - | + | + |
| Succinate | + | - | - |
| Ethanol | - | n.r. | n.r. |
| Methanol | - | n.r. | n.r. |
| **Electron acceptor** | | | |
| Nitrate | - | + | - |
| Sulfate | - | - | - |
| Thiosulfate | + | + | + |
| Fumarate | - | - | - |
| Perchlorate | + | n.r. | n.r. |
| Fe (III) | - | - | n.r. |
